# Early reaction time variability predicts implicit statistical learning: a comparison of four variability indices

**DOI:** 10.64898/2026.05.29.728728

**Authors:** Emanuele Ciardo, Andreas Alexandersen, Emanuele Galladini, Dicle Karacadağ, Teodóra Vékony, Dezső Németh

## Abstract

High intra-individual reaction time variability (RTV) is traditionally viewed through a deficit perspective and interpreted as a maladaptive signature of attentional lapses, cognitive inefficiency, and systemic noise. However, theories from motor learning and the competitive neurocognitive networks framework suggest that behavioral variability and reduced top-down control might actually facilitate certain forms of implicit skill acquisition. The present study addresses the apparent conflict between these perspectives by investigating whether elevated RTV serves as an adaptive, functional precursor to implicit statistical learning. Across two independent studies, participants completed the Alternating Serial Reaction Time (ASRT) task. We quantified early RTV during the initial task phase using multiple metrics — coefficient of variation, inter-trial RTV, and ex-Gaussian parameters Sigma and Tau— to predict subsequent statistical learning. Analyses controlled for baseline response speed and early learning artifacts, and test-retest reliability measures were also evaluated. Our results show that early RTV predicted later statistical learning measured via reaction times. This predictive relationship was most consistent for metrics capturing dynamic, moment-to-moment fluctuations (inter-trial RTV and Sigma) rather than extreme attentional lapses (Tau). While the effect size was relatively small, the association remained significant after controlling for potential statistical confounds. Furthermore, early RTV demonstrated strong test-retest stability. These findings challenge the exclusively deficit-oriented perspective on behavioral noise. Instead, we propose that elevated RTV may reflect an adaptive, exploratory processing tendency, analogous to kinematic exploration in motor learning, that could support the brain’s ability to implicitly extract and model probabilistic environmental regularities.

## 1. Introduction

Whether learning a musical instrument, mastering a fast-paced video game, or acquiring an everyday skill, people differ in how quickly and efficiently they adapt to novel regularities. Such individual differences in learning are a robust feature of both visuomotor and cognitive performance, motivating efforts to identify early markers of later learning (Abeles et al., 2023). One candidate behavioural marker is trial-to-trial variability in performance, including fluctuations in reaction time (RT). Traditionally, such variability has been viewed as a byproduct of noise that the sensorimotor system must overcome to achieve stable behavior (He et al., 2016; Wu et al., 2014). More recent work, however, has challenged this view, suggesting instead that variability may, under some conditions, support learning by promoting exploration (Wu et al., 2014). At the same time, existing findings indicate that the relation between variability and learning is not uniform but depends on the task and the source of variability (He et al., 2016). This raises a fundamental question: when individuals first encounter a new regularity structure, does early performance variability simply reflect inefficiency, or can it reveal something meaningful about their later capacity to learn?

To address this question, the present study focuses on trial-to-trial fluctuations in response speed, or reaction time variability (RTV). RTV is typically extracted from continuous performance paradigms or choice RT tasks, where intra-individual fluctuations in response speed are tracked across tens or hundreds of trials. Traditionally, behavioral variability was captured using simple summary statistics like the standard deviation. However, empirical RT distributions are strictly bounded at zero and naturally right-skewed; thus, the standard deviation is often artificially inflated by outliers and fails to capture the true shape of the data (Whelan, 2008). Basic standardizations partially address this issue: the coefficient of variation (CV) controls for the overall processing speed, while the inter-trial reaction time variability (ITRV) captures the fluctuation between subsequent trials. Yet, a more principled approach involves distribution-sensitive techniques like ex-Gaussian modeling (Heathcote et al., 1991; Matzke & Wagenmakers, 2009). By convolving a normal and an exponential distribution, this method mathematically parses variability into two distinct components: Sigma, which reflects the symmetric spread of routine motor and cognitive execution, and Tau, which quantifies the stretched rightward tail of abnormally slow responses typically linked to attentional lapses or mind wandering (Leth-Steensen et al., 2000). Relying on a single performance variability measure might miss important information: each RTV index is especially sensitive to a different type of variability, as illustrated in **Figure 1**. In the present study, we tested four complementary approaches to quantifying RTV—CV, ITRV, Sigma and Tau—to examine their predictive relation to later learning and to evaluate their reliability.

**Figure 1.**
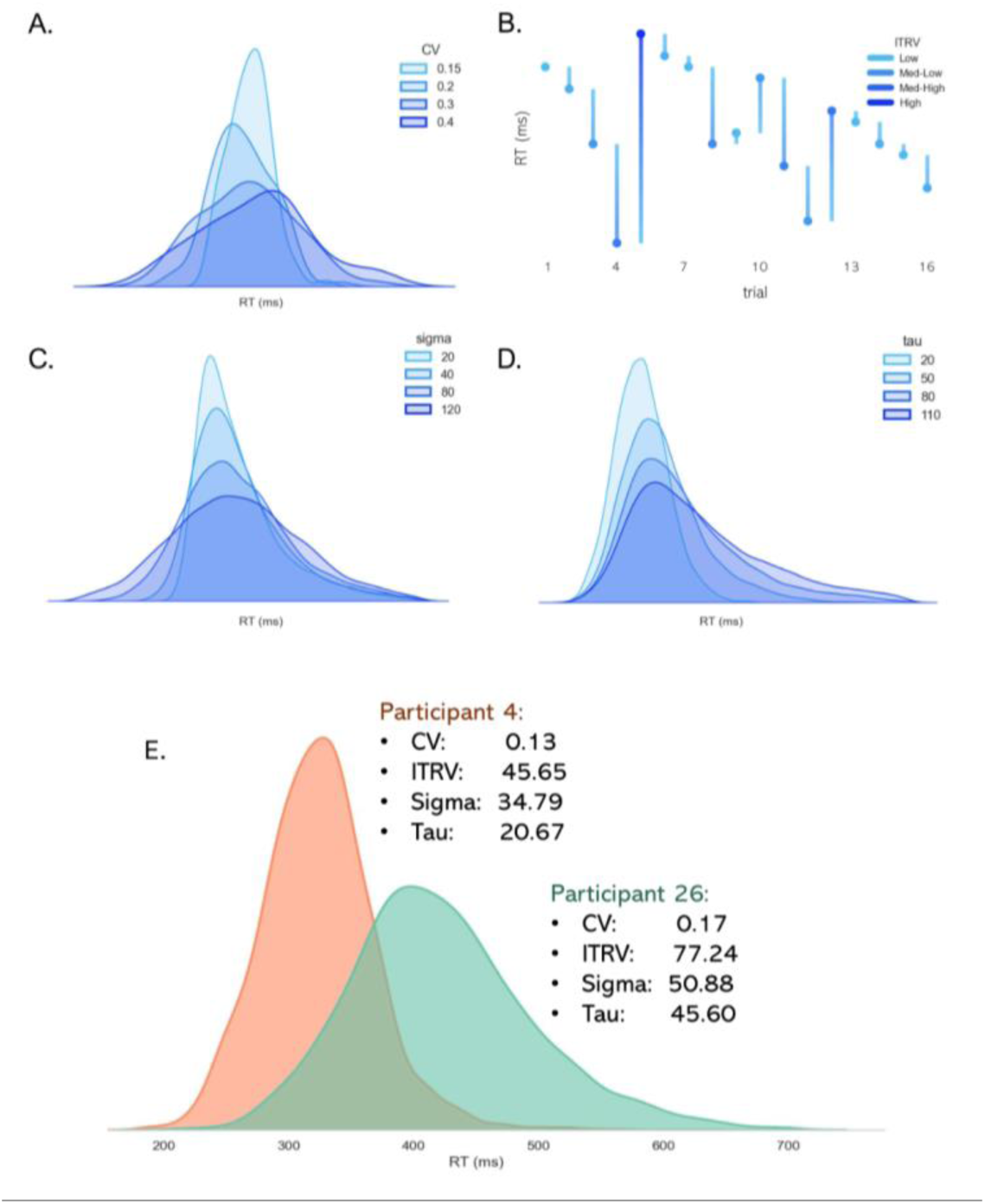
Illustration of the RTV metrics. **(A-D)** Simulated data illustrating how variations in each specific RTV metric manifest visually. **(A)** Increasing the CV proportionally widens the overall spread of the distribution relative to the mean. **(B)** ITRV captures moment-to-moment stability, depicted here by the absolute differences in RT across consecutive trials (where color depth indicates the magnitude of the trial-to-trial fluctuation). **(C)** Increasing the ex-Gaussian parameter Sigma symmetrically broadens the normal component of the distribution. **(D)** Increasing the ex-Gaussian parameter Tau specifically elongates the positive tail of the distribution, modeling an increase in abnormally slow responses. **(E)** Empirical RT distributions for two representative participants, highlighting distinct profiles across the four evaluated RTV measures. RTV = Reaction time variability; CV = Coefficient of variation; ITRV = Inter-trial RTV; SL = Statistical learning.

While the cognitive control literature often frames behavioral variability as a deficit, findings from the related field of visuomotor learning suggest it may also serve an adaptive function. However, it is important to distinguish the chronometric fluctuations (i.e., when a response is initiated), captured by RTV, from the kinematic fluctuations (i.e., how a movement is executed). In the motor domain, the seminal work of Wu et al. (2014) demonstrated that higher baseline kinematic variability predicted faster subsequent learning rates, conceptualizing this noise as a form of active exploration. This finding was later refined and partly challenged, showing that the relationship is task-specific (He et al., 2016); execution noise only predicts faster learning when it heightens uncertainty in a way that forces greater reliance on corrective feedback. These findings suggest that while behavioral variability can facilitate learning, it is not a monolithic construct, and different sources of variability yield divergent learning outcomes. Transitioning from kinematic to chronometric measures, research directly linking RTV to statistical learning remains scarce, and existing findings are mixed. For instance, applying machine learning to early behavioral features during a finger-tapping task—utilizing metrics akin to those in the RTV literature— failed to predict end-of-task learning from early behavior (Abeles et al., 2023). However, the explicit nature of the finger-tapping task and the broad search space of the algorithm adopted limit the generalizability to implicit SL. Other work has shown more promising associations. For instance, it has been shown in a Serial Reaction Time (SRT) task utilizing a long, 32-element sequence that higher RTV was associated with better sequence-specific learning, indexed by faster responses to sequence elements compared to random stimuli (Verstynen et al., 2012). While this shows the potential utility of RTV, the design could not address its predictive power for implicit SL: firstly, because the SRT task is rarely entirely implicit, and secondly because the relationship between variability and learning was only observed correlationally within the same block. A temporally separated design is required to determine whether early RTV functions as a precursor of implicit SL.

An important step toward addressing this question came from a study examining individual differences in implicit SL using the ASRT task (Stark-Inbar et al., 2017). In this paradigm, participants learn non-adjacent, second-order probabilistic dependencies where some stimulus triplets occur more frequently than others, leading to faster responses for high-probability stimuli without explicit awareness or reinforcement (Stark-Inbar et al., 2017; Vékony et al., 2022). The authors found that a greater CV, computed from blocks 2 and 3—excluding the initial familiarization block—was associated with better average SL across later blocks (blocks 4–45). This provided important initial evidence that behavioral variability is related to learning in this task, yet several questions remain unresolved. First, the study relied on a highly limited sample size, with only 21 participants included in the final correlational analyses, restricting statistical power and generalizability. Second, the learning measure was an average across almost the entire task, which is theoretically different from predicting the final learning outcome by the end of the task. Thirdly, RTV was operationalized using only a single summary measure, which could not address whether the predictive power stems from dynamic, moment-to-moment fluctuations or from extreme attentional lapses. Finally, establishing a robust predictive relationship requires control for statistical confounds—such as the inherent correlation between baseline response speed and variability, or the artifacts of early learning itself—which were beyond the scope of the original design. To address these limitations, the present study utilizes a comprehensive, well-powered approach to examine whether multiple, carefully isolated measures of early RTV predict the final level of implicit SL.

Building on this prior work, the present study pursued three primary objectives to clarify the role of early behavioral variability in implicit statistical learning. First, we aimed to validate multiple RTV indices (namely CV, ITRV, Sigma, and Tau) by evaluating their reliability and suitability as individual difference measures within the ASRT paradigm. Second, we tested whether RTV measured during the earliest phase of sequence exposure predicts the final level of SL achieved by the end of the task. To capture this critical window, we specifically focused on the first five sequence blocks (from a total of 25). This specific timeframe was chosen because it represents the period participants first encounter the structured input and actively begin to extract probabilistic regularities from the stream of stimuli. Third, to determine the temporal boundary of this relationship, we assessed whether the predictive power of early RTV over later learning extends even earlier in the experiment—specifically, to RTV calculated during the initial random training blocks, when participants were still familiarizing themselves with the basic stimulus-response mechanics. Ultimately, this multi-metric approach allows us to determine whether the link between early RTV and later learning depends on how variability is operationalized, and whether it reflects a general task-familiarization state or the active onset of structure extraction.

## 2 Materials and Methods

### 2.1 Study 1

#### 2.1.1 Participants

One hundred eighty-nine healthy young adults (109 women, 80 men; mean age = 22.05 years, range = 18–31; mean education = 15.15 years, range = 12–23) were recruited through online advertisements. Inclusion criteria were right-handedness, age under 35 years, and no or limited musical training (defined as <10 years of practice). Participants reported no current neurological or psychiatric conditions and no use of psychoactive medication. All participants provided written informed consent and received financial compensation. The Comité de Protection des Personnes approved the study (CPP Est I; ID: RCB 2019-A02510-57). While the full experimental protocol spanned three testing days, the present study focuses exclusively on Day 1 for the main analyses and Day 3 for the test-retest reliability assessment (see **Supplementary Methods** for further details).

#### 2.1.2 Task and procedure

Implicit statistical learning was assessed using a modified version of the ASRT task (Howard & Howard, 1997; Pedraza et al., 2025) implemented in Presentation software (Neurobehavioural Systems, Inc). Refer to **Figure 2 (A-C)** for a visualization of the task. On each trial, a yellow arrow appeared centrally for 200 ms and pointed in one of four directions (left, up, down, right). Participants were instructed to press the key corresponding to the direction of the arrow as quickly and accurately as they could. The arrow was immediately followed by a fixation cross for 500 ms. Participants were allowed to respond for 700 ms from the onset of the arrow stimulus. Responses were made using a four-button Cedrus RB-530 response pad (Cedrus Corporation, San Pedro, CA). Finger placement was standardized: up = left index, left = left thumb, right = right index, down = right thumb. Feedback was provided for incorrect or missing responses. Following a correct response, the fixation cross remained on screen for an additional 750 ms. Incorrect responses triggered an “X” for 500 ms, and omitted responses triggered an exclamation mark (“!”) for 500 ms; in both cases, feedback was followed by a 250 ms fixation cross before the next trial.

**Figure 2.**
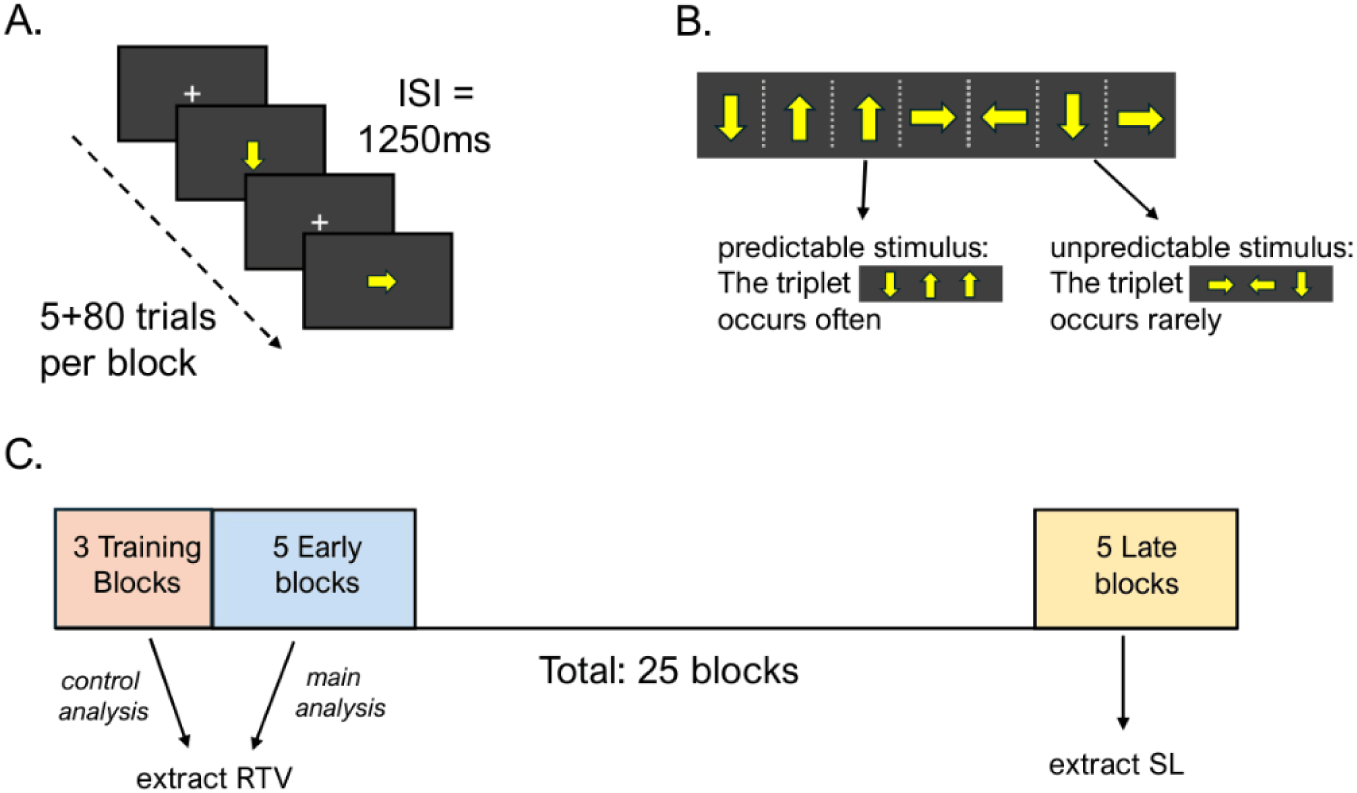
Experimental design. **(A)** Schematic of the task. Each block comprised 85 trials: 5 initial random stimuli followed by 80 alternating probabilistic stimuli. On each trial, a yellow arrow pointing in one of four directions (up, down, left, or right) appeared for 200 ms. This was followed by a 1250 ms inter-stimulus interval (ISI) featuring a fixation cross. Participants were given 700 ms from stimulus onset to respond via the corresponding key. **(B)** Illustration of predictable versus unpredictable stimuli based on triplet transition probabilities. Because pattern elements alternate with random elements, certain three-element combinations (triplets) occur more frequently, making their final elements more predictable. **(C)** Experimental block design. The session began with 3 sequence-free training blocks to habituate participants, followed by 25 sequence-embedded main blocks (30 blocks in Study 2). For the primary analyses, early RTV was extracted from the first 5 sequence blocks; a separate control analysis utilized RTV from the initial training blocks. Statistical learning, indexed by faster reaction times to predictable stimuli, was quantified exclusively during the late task phase (blocks 21 to 25). RTV = Reaction time variability.

Unbeknownst to participants, stimuli followed an alternating probabilistic structure in which pattern (P) elements alternated with random (R) elements (e.g., 4–R–2–R–3–R–1–R, where numbers indicate fixed locations/directions and R indicates a random element). This alternating structure yields unequal occurrence probabilities for runs of triplets (three consecutive trials): some triplets occur frequently (high-probability), whereas others occur infrequently (low-probability). High-probability triplets can arise in both P–R–P and R–P–R configurations, whereas low-probability triplets can arise only in R–P–R configurations, making high-probability triplets five times more frequent. SL in the ASRT task is typically indexed by faster responses to the final element of high-probability triplets (predictable stimuli) compared with the final element of low-probability triplets (unpredictable stimuli), without participants acquiring explicit awareness of the sequence structure.

Before the main task, participants completed an initial training phase designed to familiarize them with the procedure. This phase consisted of three training blocks of 85 trials containing entirely random stimuli with no underlying sequence. During this time, participants were explicitly notified that they were completing practice trials and that the main task would commence afterward. Following this training, the task began, consisting of 25 blocks, each containing 85 stimuli: five initial randomly ordered stimuli (warm-up) followed by 10 repetitions of an eight-element alternating sequence. Upon completion of the experiment, a verbal interview was conducted to assess whether participants had acquired explicit awareness of the underlying alternating sequence.

#### 2.1.3 Data analysis

Our analyses focused on the relationship between early RTV and later SL. To ensure a strict temporal separation between the predictor and outcome variables, SL was quantified exclusively during the late phase of the task (blocks 21–25). In contrast, early RTV was evaluated in two distinct ways: either during the first five sequence-embedded blocks or, in a separate analysis, during the three sequence-free training blocks. This temporal isolation allowed us to test whether initial individual differences in variability predict subsequent learning, effectively eliminating any measurement overlap between the two metrics.

##### 2.1.3.1 Data preprocessing

Before computing the behavioral metrics, raw RT data underwent distinct filtering procedures tailored to the requirements of the RTV and SL analyses.

For the RTV analyses the objective was to preserve the full empirical shape of the distribution. Therefore, the only criterion was the exclusion of error trials, leaving all correct responses (hits) for the calculation of the metrics. We did not include a cutoff for abnormally slow answers because the maximum response time was 700 ms.

The preprocessing pipeline for SL was stricter. Firstly, data were restricted to the late phase of the task (blocks 21–25). To ensure that responses reflected genuine sequence-specific SL rather than task familiarization or low-level motor artifacts, we excluded sequence-free warm-up trials, incorrect responses, trills, and repetitions. Specifically, trills (alternating sequences such as 1-2-1) and repetitions (repeated elements such as 1-1-1) were removed because they typically elicit artificially faster RTs driven by basic visuomotor facilitation. Retaining these structural artifacts could artificially inflate performance and confound the measurement of true probabilistic learning (Song et al., 2007). Additionally, extremely fast, anticipatory responses (RT < 50 ms) were discarded. Finally, to eliminate intra-individual RT outliers without assuming a normal data distribution, we applied a Median Absolute Deviation (MAD) approach (Leys et al., 2013), calculated independently for each participant. We computed the robust standard deviation as 1.4826 × MAD. Any trials falling outside the participant-specific boundaries of the median ± 3 robust standard deviations were excluded from the final SL calculations.

##### 2.1.3.2 Quantification of reaction time variability

Early RTV was estimated separately for each participant using four indices: CV, ITRV, and the ex-Gaussian parameters Sigma and Tau. See **Figure 1(A-E)** for a visualization of the indices and how they are differentially sensitive to RT distributions and response patterns. All calculations were performed at the block level, producing a single RTV score per block for each participant.

The CV was calculated to control for individual differences in overall processing speed, defined as the standard deviation of RTs divided by the mean RT:

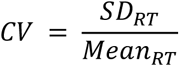

ITRV was computed to capture moment-to-moment fluctuations, defined as the mean absolute difference in RT between consecutive trials. For a block of *N* trials:

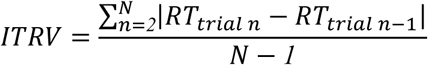

Furthermore, we fitted ex-Gaussian models to each participant’s early RT distribution using the *exponnorm.fit* function from the *scipy.stats* library in Python (Virtanen et al., 2020); the optimization algorithm successfully converged for 100% of the block distributions. This method mathematically parses variability into two distinct components by modeling the empirical RT distribution as the convolution (sum) of a normal distribution and an exponential distribution. In this model, Sigma represents the standard deviation of the symmetric Gaussian component. Conversely, Tau represents the mean of the exponential component, capturing the stretched tail of the distribution typically associated with abnormally slow responses. Based on the implementation in *scipy.stats*, if *K* is the fitted shape parameter and *scale* is the fitted scale parameter, these metrics are derived as follows:

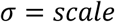

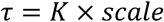

Finally, to ensure that highly abnormal performance blocks did not skew the aggregate variability estimates, we applied a standard Z-score exclusion criterion to the newly computed metrics. Any block featuring an RTV score greater than 3 standard deviations from the overall metric mean was removed from subsequent analyses. Applying this criterion resulted in minimal data loss: For the first 5 ASRT blocks, out of 945 total blocks (189 participants × 5 blocks), this procedure excluded 2 blocks for CV, 6 for ITRV, 15 for Sigma, and 2 for Tau. Similarly, during the 3 training blocks (189 participants × 3 blocks = 567), 2 blocks were excluded for CV, 2 for ITRV, 10 for Sigma, and 1 for Tau. This step led to the exclusion of 1 participant, whose 3 blocks all exceeded the 3 standard deviations threshold.

##### 2.1.3.3 Quantification of statistical learning

Statistical learning performance was quantified using data from blocks 21 through 25. Following a RT-based approach, SL was computed as the difference in mean RT between unpredictable (low-probability) and predictable (high-probability) stimuli:

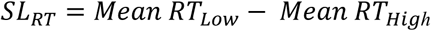

##### 2.1.3.4 Main regression analyses

To determine whether early behavioral variability predicts later SL, we conducted a series of simple linear regression analyses. The dependent variable was SL_RT_, and the independent variable was one of the four early RTV metrics (CV, ITRV, Sigma, or Tau).

##### 2.1.3.5 Control analyses: Isolating sequence-free variability

A potential confounding factor in our primary analyses is that early RTV estimates could be artificially inflated by the rapid onset of SL itself. Specifically, if participants begin responding faster to high-probability triplets during the early phase of the task, combining both predictable and unpredictable trial types could widen the overall RT distribution and artificially inflate RTV.

To ensure our metrics captured genuine, trait-like variability rather than early learning artifacts, we conducted control analyses using data exclusively from the initial three training blocks. Because these training trials consisted of entirely random, sequence-free stimuli, the resulting RTV calculations could not be confounded by emerging SL. Following the same analytical approach as the main models, the four sequence-free RTV metrics (CV, ITRV, Sigma, and Tau) were computed, and separate simple linear regressions were conducted to assess how well they predicted SL_RT_ by the end of the task.

##### 2.1.3.6 Control analysis: Accounting for baseline processing speed

Because indices of intra-individual variability often scale proportionally with overall response speed, it is important to verify that the relationship between RTV and SL cannot be explained simply by individual differences in baseline processing speed. To test this, we conducted multiple linear regression analyses predicting SL_RT_, entering both the specific early RTV metric (CV, ITRV, Sigma, or Tau) and mean RT simultaneously into the model. To ensure the reliability of the coefficient estimates and to confirm that the models were not compromised by shared variance between mean RT and the RTV metrics, we calculated Variance Inflation Factors (VIF) for all predictors.

##### 2.1.3.7 Stability analysis

To assess the test–retest reliability of the RTV measures, we leveraged the study’s longitudinal design, which included a follow-up testing session two days later. Out of the 189 recruited participants, 8 did not complete the follow-up session, reducing the sample size for this analysis to 181. For mean RT and each RTV metric (CV, ITRV, Sigma, and Tau), stability was quantified using Intraclass Correlation Coefficients (ICC; Koo & Li, 2016). We calculated both absolute agreement and consistency to provide an estimate of session-to-session reliability.

To evaluate how these reliability estimates depended on the amount of data available, we performed a cumulative stability analysis. We recomputed each metric and its corresponding ICC at each step while progressively increasing the number of task blocks included in the calculation (from 1 to 25 blocks).

### 2.2 Study 2

#### 2.2.1 Participants

To verify our primary findings in an independent sample, Study 2 utilized a dataset originally collected by Simor and colleagues (2025). The sample consisted of 36 university students (mean age = 22.1 years, SD = 1.27, 30 female) who participated in exchange for partial course credit. Participants reported no history of psychiatric, neurological, or chronic somatic disorders; were not taking medications known to affect alertness, mood, cognition, or sleep; and had normal or corrected-to-normal vision.

Right-handedness was confirmed using the Edinburgh Handedness Inventory (Oldfield, 1971). Participants were naive to the experiment’s purpose before the session and were fully debriefed afterward, at which point verbal reports were collected to confirm they remained naive to the sequence structure. All participants provided written informed consent, and the study protocol was approved by the Research Ethics Committee of Eötvös Loránd University.

#### 2.2.2 Task and procedure

The modified ASRT task and the overall experimental procedure were identical to those described in Study 1, with one exception: the main task in Study 2 comprised 30 blocks rather than 25.

#### 2.2.3 Data preprocessing and analysis

Data preprocessing, trial-level filtering, and the quantification of both early RTV and late Statistical Learning (SL_RT_) followed the same protocols established in Study 1. To maintain analytical consistency across the two studies, SL performance in Study 2 was quantified using the identical temporal window (blocks 21 through 25), even though the participants completed 30 blocks in total.

As in the first study, we applied a standard Z-score exclusion criterion (> 3 SD from the overall metric mean) to remove highly abnormal performance blocks before aggregating the early RTV estimates. This procedure resulted in negligible data loss: During the first 5 ASRT blocks, out of 180 total blocks (36 participants × 5 blocks = 180 blocks), no blocks were excluded for CV and ITRV, 1 block was excluded for Sigma, and 1 block was excluded for Tau. Regarding the 3 random (training) blocks, out of 108 total blocks (36 participants × 3 blocks = 108 blocks), no blocks were excluded for CV and Tau, 1 block was excluded for Sigma, and 1 block was excluded for ITRV.

#### 2.2.4 Statistical analyses

The statistical analyses for Study 2 directly mirrored the primary and control models conducted in Study 1, except for the test-retest stability analysis (which was specific to the multi-day design of Study 1). Specifically, we evaluated the following models: 1) Main regression analyses: Simple linear regressions predicting end-of-task SL from the four early RTV metrics (CV, ITRV, Sigma, and Tau) calculated over blocks 1–5. 2) Control analyses isolating sequence-free variability: Simple linear regressions predicting SL using RTV estimates derived exclusively from the initial, sequence-free random training blocks. 3) Control analyses accounting for baseline speed: Multiple linear regressions predicting SL by entering the early RTV metric and mean RT simultaneously. This ensured that the observed relationships could not be explained solely by individual differences in baseline processing speed.

## 3. Results

### 3.1 Study 1

#### 3.1.1 Inter-correlation and test-retest reliability of early RTV metrics

As an initial exploration of the relationship between the extracted measures, we computed a Pearson correlation matrix linking early (block 1 to 5) RTV metrics and processing speed (mean RT), and subsequent statistical learning (SL_RT_). These relationships, corrected for multiple comparisons using false discovery rate (*q* < .05), are visualized in **Figure 3A**. Overall, RTV measures were highly correlated with each other (all *r* ≥ .58; all p < .001), except for the pair Tau and Sigma (*p* = .10; *r* = .12). Sigma and ITRV presented a high correlation with mean RT (*r* ≥.57, *p* < .001), while the correlation was weaker but significant for CV and Tau (both *r* = .21, *p* < .01). Importantly, SL_RT_ was weakly but significantly correlated with CV, ITRV and Sigma (*r* ≥ .22, *p* < .01). In contrast, SLRT was not correlated with Tau or mean RT. Furthermore, as illustrated in **Supplementary Figure 3**, mapping the normalized RTV measures across the task blocks reveals a clear, unified negative trend over time for all four indices.

**Figure 3.**
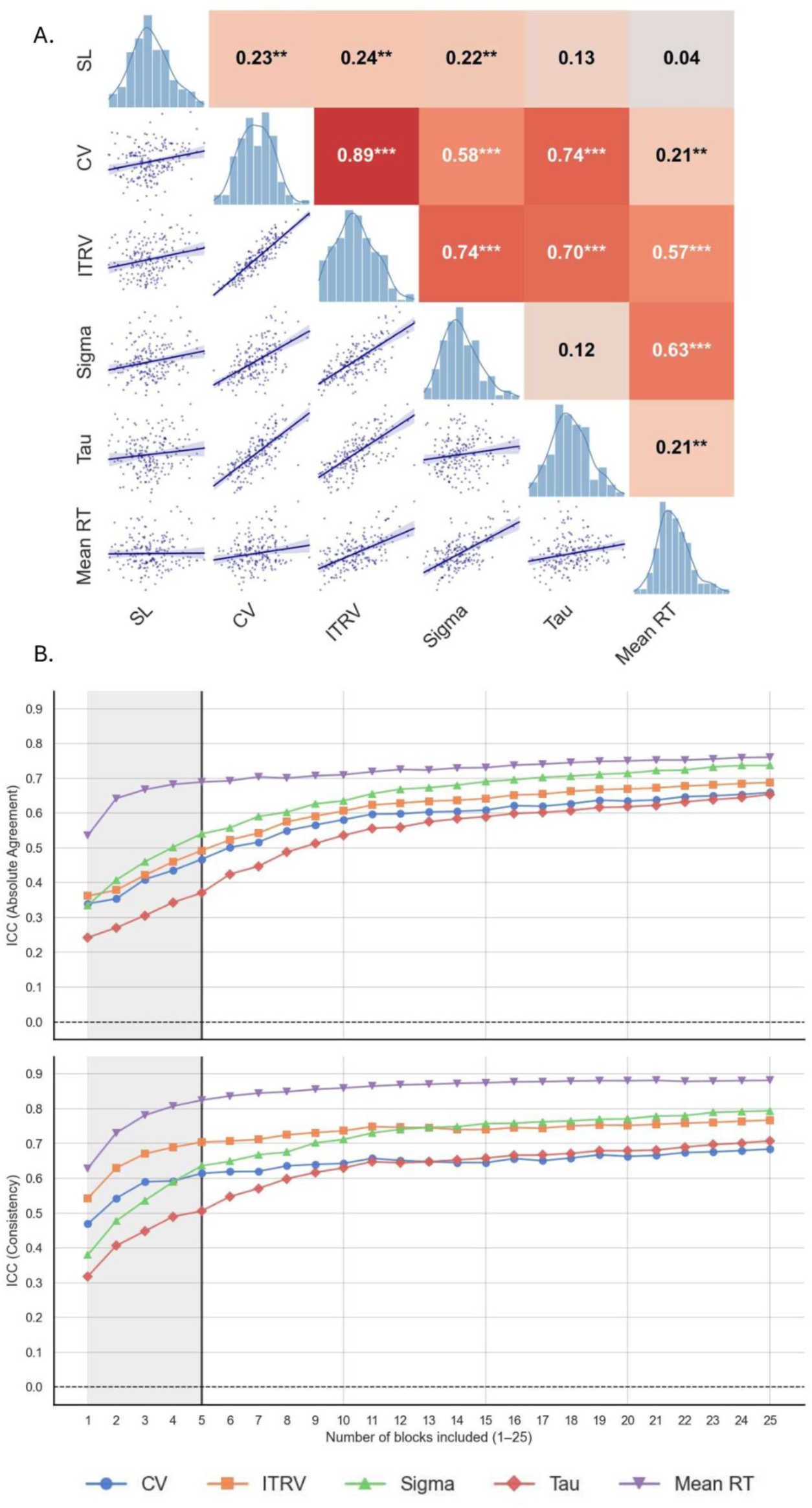
Descriptive analysis on RTV. **(A**) Pearson correlation matrix of early (blocks 1 to 5) RTV and mean RT and implicit SL in late blocks (21 to 25). The evaluated RTV metrics include the CV, ITRV, and the ex-Gaussian parameters Sigma and Tau. SL_RT_ reflects the RT difference between low- and high-probability triplets at the end of the task. Cell shading indicates the magnitude and direction of associations; associations were all positive. (**B)** Test-retest reliability of RTV metrics as a function of task length. The plots display Intraclass Correlation Coefficients (ICC) for Absolute Agreement (top panel) and Consistency (bottom panel) calculated cumulatively across 25 blocks of the ASRT task. Metrics include mean RT (purple), Sigma (green), ITRV (orange), CV (blue), and Tau (red). The shaded area highlights the first five blocks, indicating the "early RTV" window used in subsequent predictive analyses. Stability for all metrics increases rapidly within the first few blocks before reaching a plateau. Horizontal dashed lines at 0.0 indicate the baseline for no correlation. **p < .01, ***p < .001.

To quantify the test–retest reliability of each index (CV, ITRV, Sigma, Tau, alongside mean RT for comparison) across the 48-hour delay, we computed Intraclass Correlation Coefficients (ICCs). We evaluated the data using two distinct ICC models (Koo & Li, 2016). First, to assess *relative consistency*—how well participants maintained their rank order relative to one another across the two sessions—we utilized a two-way mixed-effects, consistency, single measurement model [ICC(3,1)]. Second, to evaluate *absolute agreement*—the degree to which participants’ exact numerical scores replicated across sessions—we employed a two-way mixed-effects, absolute agreement, single measurement model [ICC(2,1)].

We first examined how the stability of these measures scaled with the volume of data included in the estimation. As illustrated in **Figure 3B**, reliability increased sharply when moving from a single block to multiple blocks before plateauing relatively quickly. This trajectory indicates that aggregating a modest amount of data yields a reliable measure of an individual’s behavioral signature. Based on the plateau in consistency, we focused our primary stability analysis on the aggregated data from the first five blocks.

Over the 48-hour delay, the variability indices extracted from these initial five blocks demonstrated moderate relative consistency. ITRV showed the highest relative stability (ICC = .70, 95% CI [.62, .77], *F*(180, 180) = 5.76, *p* < .001), followed by Sigma (ICC = .64, 95% CI [.54, .72], *F*(180, 180) = 4.49, *p* < .001), CV (ICC = .61, 95% CI [.51, .70], *F*(180, 180) = 4.19, *p* < .001), and Tau (ICC = .51, 95% CI [.39, .61], *F*(180, 180) = 3.05, *p* < .001). As anticipated absolute agreement was lower than relative consistency—likely reflecting systematic practice or habituation effects that shifted overall response times between sessions. Nevertheless, absolute agreement across these first five blocks remained notable given the nature of the data, ranging from moderate for Sigma (ICC = .54, 95% CI [.21, .72]), ITRV (ICC = .49, 95% CI [-.06, .76]), and CV (ICC = .47, 95% CI [.04, .70]), to poor for Tau (ICC = .37, 95% CI [.02, .60]). Finally, as expected, mean RT extracted from the same five-block window demonstrated the highest test-retest reliability, showing good relative consistency (ICC = .82, 95% CI [.77, .87], *F*(180, 180) = 10.41, *p* < .001) and moderate absolute agreement (ICC = .69, 95% CI [.09, .87]).

Taken together, these results indicate that while the exact numerical values of early RTV underwent a general shift across sessions, the relative ranking between participants remained robust, confirming early RTV as a sufficiently stable individual-difference metric in the ASRT task.

#### 3.1.2 Does early RTV predict later SL?

Before moving to the main analyses, we confirmed that the participants exhibited statistical learning in the late blocks of interest, from 21 to 25. A two-tailed, one-sample t-test showed the presence of SL_RT_ (*M* = 4.73, *SD* = 6.67, *t*(188) = 9.75, *p* < .0001, *d* = 0.71). Importantly, post-experiment verbal reports confirmed that no participants reached explicit awareness of the sequence structure, verifying the implicit nature of this learning.

To establish whether measures of RTV during the early task blocks predicted SL_RT_ by the end of the task, a series of simple linear regression analyses was conducted. As can be seen in **Figure 4A**, three of the four early RTV metrics emerged as significant positive predictors: CV (*F*(1, 187) = 10.37, *p* = .002, *R*² = .053, *b* = 79.08, *β =* 0.23, *SE* = 24.56, *t* = 3.22), ITRV (*F*(1, 187) = 10.97, *p* = .001, *R*² = .055, *b* = 0.15, *β =* 0.24, *SE* = 0.04, *t* = 3.31), and Sigma (*F*(1, 187) = 9.54, *p* = .002, *R*² = .049, *b* = 0.19, *β =* 0.22, *SE* = 0.06, *t* = 3.09). Tau did not reach significance, showing only a marginal positive trend (*F*(1, 187) = 3.41, *p* = .066, *R*² = .018, *b* = 0.08, *β =* 0.13, *SE* = 0.04, *t* = 1.85). Therefore, various individual measures of RTV in the early stages of the task predicted later SL

**Figure 4.**
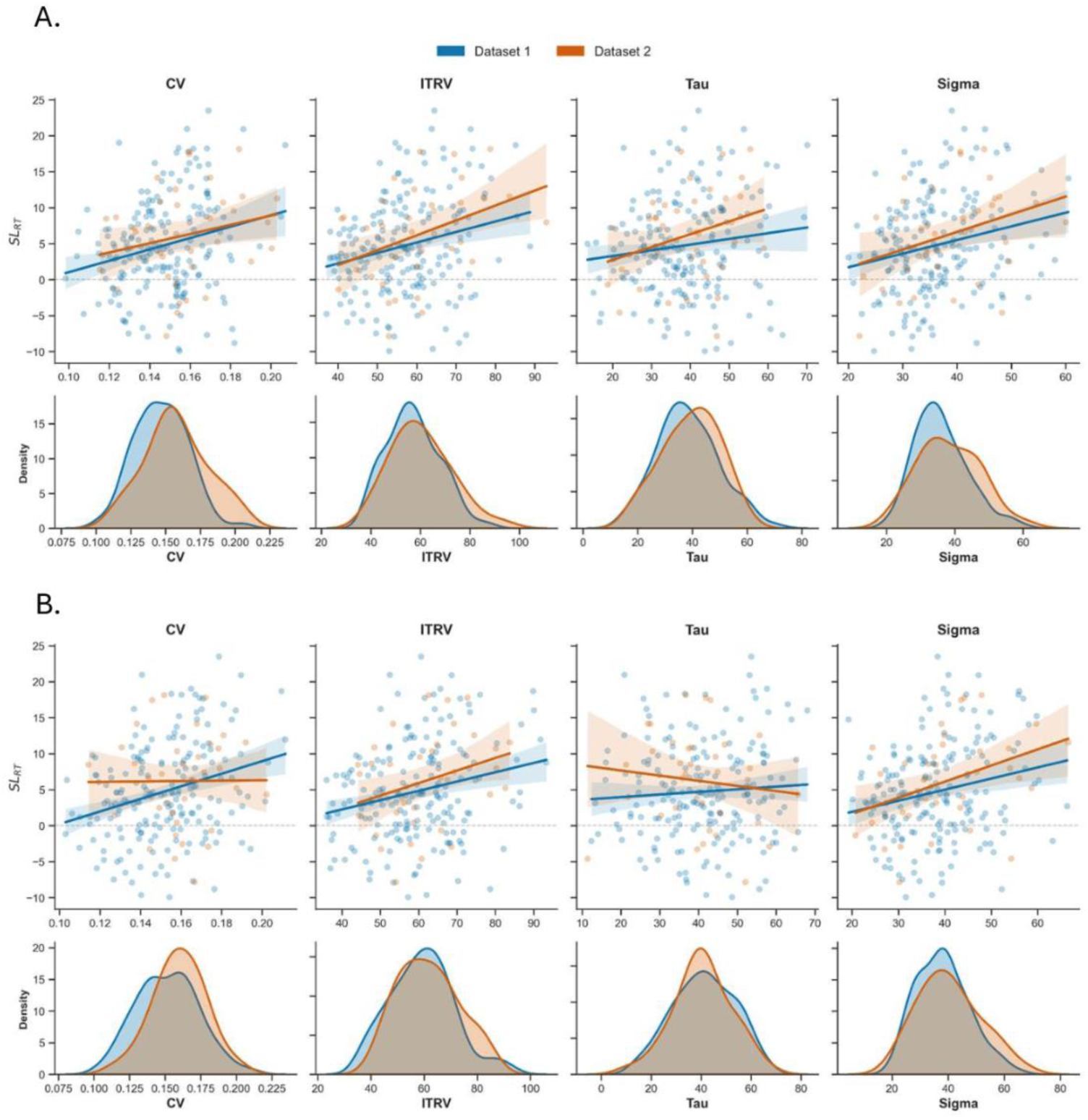
Overview of the relationship between four early RTV metrics and end-of-task statistical learning. Data from Study 1 is shown in blue, and Study 2 is shown in orange. **(A)** RTV calculated during early sequence blocks (blocks 1–5). **Upper row**: Scatterplots and linear regression fits (solid lines) with 95% confidence intervals (shaded bands) predicting SL_RT_. The plots show upward slopes for CV, ITRV, and Sigma. ITRV and Sigma are statistically significant positive predictors in both independent datasets, while CV is significant only in Study 1. In contrast, Tau displays a relatively flat, non-significant trend across both studies. **Bottom row**: Density plots of the RTV distributions. **(B)** RTV calculated during sequence-free training blocks. **Upper row:** Illustrates the same predictive relationships, but using RTV extracted before any probabilistic sequence was introduced. Sigma remains a significant positive predictor of SL_RT_ across both datasets, even in the absence of a sequence.

#### 3.1.3 Do early RTV effects persist in the absence of sequence structure?

To rule out the possibility that early RTV was artificially inflated by the onset of SL itself (i.e., variance increasing simply because participants were beginning to respond faster to predictable versus unpredictable trials), we repeated the regression analyses using RTV computed exclusively from the initial training blocks, which consisted solely of random stimuli without any underlying sequence. Restricting the data to this initial window resulted in the exclusion of one participant from the Sigma analysis, as all three of their random blocks were removed because they diverged more than 3 SD from the mean.

The overall pattern of results remained similar, with the same RTV metrics reaching statistical significance. Results are summarized in **figure 4B**. Early variability continued to significantly predict SL_RT_ when using CV (*F*(1, 187) = 16.69, *p* < .001, *R*² = .082, *b* = 87.35, *β =* 0.29, *SE* = 21.38, *t* = 4.08), ITRV (*F*(1, 187) = 10.92, *p* = .001, *R*² = .055, *b* = 0.13, *β =* 0.23, *SE* = 0.04, *t* = 3.31) and Sigma (*F*(1, 186) = 9.28, *p* = .003, *R*² = .048, *b* = 0.15, *β =* 0.22, *SE* = 0.05, *t* = 3.05), while it was not predicted by Tau (*F*(1, 187) = 0.85, *p* = .356, *R*² = .005, *b* = 0.04, *β =* 0.07, *SE* = 0.04, *t* = 0.92).

Taken together, these findings demonstrate that the predictive relationship between early RTV and later SL is not simply an artifact of early sequence-induced fluctuations, but rather reflects genuine correlation between the studied measures. **Supplementary analyses** extend this result, showing that RTV remained a significant predictor of late day 1 SL even when calculated in either of the subsequent testing days.

#### 3.1.4 Controlling for mean RT

Since RTV showed high correlation with mean RT - as shown in the previous section 3.1.1 -, it is important to verify that the predictive power of RTV over SL is not driven by the mediating effect of mean RT. To test this, we conducted multiple linear regression analyses predicting SL_RT_, with both the specific RTV metric and mean RT entered simultaneously as predictors. Variance Inflation Factors (VIF) for all models were below 1.7, confirming that the models were not compromised by multicollinearity.

The predictive power of early variability remained highly significant. Controlling for mean RT, significant unique variance in SL_RT_ was still explained by CV (*F*(2, 186) = 5.18, *p* = .007, *R*² = .05, *b* = 78.21, *β =* 0.23, SE = 25.00, *t* = 3.13, *p(t)* = .002), ITRV (*F*(2, 186) = 6.23, *p* = .002, *R*² = .06, *b* = 0.18, *β =* 0.29, SE = 0.05, *t* = 3.45, *p(t)* < .001) and Sigma (*F*(2, 186) = 5.89, *p* = .003, *R*² = .06, *b* = 0.26, *β =* 0.30, SE = 0.08, *t* = 3.35, *p(t)* = .001). As in the primary analyses, Tau did not emerge as a significant predictor (*F*(2, 186) = 1.73, *p* = .180, *R*² = .02, *b* = 0.07, *β =* 0.13, SE = 0.04, *t* = 1.71, *p(t)* = .090). In all models, mean RT itself was not a significant predictor of learning outcomes (all *p* > .140).

### 3.2 Study 2

#### 3.2.1 Does early RTV predict later SL?

To validate the findings from our primary study, we sought to replicate the predictive relationship between early RTV and later SL_RT_ in an independent cohort. Firstly, we again confirmed with a 2-tailed, one-sample t-test that the participants extracted the probabilistic sequence by performing faster in predictable trials (*M* = 5.13, *SD* = 4.43, *t*(35) = 6.95, *p* < .0001, *d* = 1.16). Post-experiment verbal reports confirmed that no participants reached explicit awareness of the sequence structure. Next, we conducted a series of simple linear regression analyses, mirroring what applied in Study 1. Specifically, both ITRV (*F*(1, 34) = 6.01, *p* = .019, *R²* = .150, *b* = 0.21, *β =* 0.39, *SE* = 0.08, *t* = 2.45) and Sigma (*F*(1, 34) = 4.38, *p* = .044, *R²* = .114, *b* = 0.25, *β =* 0.34, *SE* = 0.12, *t* = 2.09) emerged as significant positive predictors of learning. Conversely, CV (*F*(1, 34) = 1.75, *p* = .20, *R²* = .049, *b* = 63.98, *β =* 0.22, *SE* = 48.49, *t* = 1.32) and Tau (*F*(1, 34) = 2.98, *p* = .09, *R²* = .081, *b* = 0.18, *β =* 0.28, *SE* = 0.10, *t* = 1.73) did not reach statistical significance in this study. The results overall match what was observed in Study 1, where early variability successfully predicted end-of-task SL_RT_, with the difference that CV was no longer a significant predictor in Study 2.

#### 3.2.2 Do early RTV effects persist in the absence of sequence structure?

Following the analytical pipeline applied in Study 1, we verified that this predictive relationship was not an artifact of early sequence-induced fluctuations. We repeated the regression analyses, extracting variability metrics exclusively from the initial random training blocks, which lacked any underlying sequence.

The results remained similar to the primary validation analysis. Sigma (*F*(1, 34) = 5.72, *p* = .022, *R²* = .144, *b* = 0.22, *β =* 0.38, *SE* = 0.09, *t* = 2.39) remained a significant positive predictor of later SL_RT_, while ITRV only reached a positive trend level (*F*(1, 34) = 3.39, *p* = .074, *R²* = .09, *b* = 0.17, *β =* 0.30, *SE* = 0.09, *t* = 1.84). As before, CV (*F*(1, 34) = 0.00, *p* = .962, *R²* = .000, *b* = 2.86, *β =* 0.008, *SE* = 60.35, *t* = 0.00) and Tau (*F*(1, 34) = 0.55, *p* = .465, *R²* = .016, *b* = -0.07, *β = -*0.13, *SE* = 0.1, *t* = -0.74) failed to reach significance. This confirms that, for the metrics sensitive to this validation sample, the observed relationship continues to reflect a genuine, sequence-independent behavioral signature rather than early learning itself.

#### 3.2.3 Controlling for mean reaction time

Finally, we evaluated whether early RTV predicted learning independently of overall processing speed by entering both the specific RTV metric and mean RT simultaneously into multiple linear regression models (all VIFs < 1.7).

In contrast to the statistically solid effects observed in the larger Study 1 sample, controlling for mean RT in Study 2 attenuated the statistical significance of the variability metrics. The model including ITRV and mean RT remained a significant (*F*(2, 33) = 3.51, *p* = .042, *R²* = .18), but the effect of ITRV was reduced to a marginal trend (*b* = 0.17, *β =* 0.32, *SE* = 0.09, *t* = 1.82, *p(t)* = .077). Sigma no longer reached significance (*F*(2, 33) = 2.70, *p* = .082, *R*² = .14, *b* = 0.18, *β =* 0.26, *SE* = 0.13, *t* = 1.37, *p(t)* = .181), and both CV and Tau remained non-significant (CV: *F*(2, 33) = 2.64, *p* = .086, *R*² = .14, *b* = 62.13, *β =* 0.21, *SE* = 46.86, *t* = 1.33, *p(t)* = .194; Tau: *F*(2, 33) = 2.64, *p* = .087, *R*² = .14, *b* = 0.14, *β =* 0.22, *SE* = 0.10, *t* = 1.32, *p(t)* = .196). This attenuation suggests that while the baseline predictive relationship replicates, isolating the unique variance of RTV from general processing speed is challenging due to the inherent shared variance between these chronometric measures.

## 4. Discussion

The present study was designed with three primary objectives: first, to validate multiple indices of reaction time variability — the coefficient of variation, the inter-trial reaction time variability, Sigma, and Tau — by evaluating their reliability and suitability as individual difference measures within a widely used statistical learning task; second, to determine whether RTV measured during the earliest phase of sequence exposure predicts the final level of statistical learning achieved; and third, to establish whether this predictive power extends to RTV calculated during the initial random training blocks. Across two independent studies, our findings demonstrated that early RTV significantly and positively predicted statistical learning in the late stage of the task, as indexed by reaction time gains for predictable stimuli, which occurred entirely outside of participants’ explicit awareness. In particular, out of the four adopted RTV metrics, the Gaussian spread Sigma and ITRV were the most consistent predictors, both of which are especially sensitive to moment-to-moment fluctuations rather than extreme lapses. Critically, this predictive relationship did not depend on the presence of the sequence itself. Early RTV predicted later statistical learning whether it was measured during the first five sequence blocks or during the three random training blocks administered before any structured input appeared. A control analysis confirmed that the effect partially survived after controlling for response speed. Furthermore, the stability analyses confirmed that these RTV measures exhibit high test-retest reliability, supporting their role as stable, trait-like markers of individual learning capacity rather than transient session-specific noise.

The idea that early behavioral noise carries information about future learning sounds intuitive, even obvious. The empirical record, however, tells a different story: this question has never been put to a direct test. Prior work on behavioral variability and learning has focused mostly on motor adaptation (He et al., 2016; Wu et al., 2014) or has utilized designs that make it difficult to isolate early variability from the ongoing learning process itself. The closest precedent comes from Stark-Inbar and colleagues (2017), who found a positive link between RT variability and SL in the same paradigm we used. Their measures were separated in time, but their learning score averaged almost the whole task. So even with that link in hand, one question stays open: did variability predict how quickly people learned, or how much they ultimately learned? Similarly, previous works described a connection between variability and sequence learning in an SRT paradigm (Verstynen et al., 2012), but because their two measures were estimated within the exact same blocks, their study could not establish whether variability came before learning or simply co-occurred with it. By estimating RTV from the absolute earliest blocks (including initial random training) and isolating learning to the final blocks of the task, the present study offers a much stricter temporal test of this relationship in implicit SL. The results show that early variability actively precedes and predicts the final magnitude of later learning, rather than just accompanying it. Our findings also differ from those of Abeles et al. (2023), who could not predict end-of-task motor skill learning from early behavior. Their task was explicit and involved finger-tapping, which likely engages different processes than implicit probabilistic sequence learning. The contrast reinforces a recurring theme in this literature: behavioral variability is not a monolithic construct. What it predicts depends on the task that elicits it and on which component of variability is measured (He et al., 2016). This is precisely why our four metrics did not behave alike, with ITRV and Sigma predicting learning while Tau did not.

Elevated RTV is traditionally interpreted as a behavioral index of reduced top-down cognitive control and fluctuating attention: within-subject evidence demonstrated that localized increases in RTV reliably precede self-reports of mind wandering (Bastian & Sackur, 2013; Zanesco et al., 2021) and consistently predict error rates in sustained attention tasks (Manly et al., 2000; Rosenberg et al., 2013). The link between variability and attentional deficits is further underscored by clinical research, where RTV has emerged as perhaps the most robust behavioral correlate of ADHD (Kofler et al., 2013). Interestingly, within the ADHD population, this elevated variability appears to be driven predominantly by rare, extreme attentional lapses—captured by the Tau parameter of the ex-Gaussian distribution—rather than by ongoing, trial-to-trial fluctuations like ITRV or Sigma (Hervey et al., 2006; Vaurio et al., 2009). Importantly, heightened RTV extends well beyond attention-specific conditions and transient mental states; it is also a recognized feature of depression, schizophrenia, and borderline personality disorder (Kaiser et al., 2008), as well as a hallmark of cognitive aging (Graveson et al., 2016; MacDonald et al., 2006). On a neurobiological level, higher RTV has been associated with compromised white matter integrity (Booth et al., 2019; Tamnes et al., 2012), reduced volume in prefrontal cortical areas (Albaugh et al., 2017), and even demonstrates a weak but significant negative correlation with general intelligence (Doebler & Scheffler, 2016). Together, these diverse lines of evidence suggest that increases in RTV are multi-determined, having various origins that range from natural fluctuations in mental states to structural decreases in neural efficiency, systemic cognitive decline, and psychiatric disorders.

However, viewed through the framework of competitive neurocognitive networks, this apparent deficit in top-down control may actually prove advantageous, as implicit learning often operates more effectively under these exact conditions. In fact, recent modeling has demonstrated that trial-to-trial chronometric fluctuations in standard cognitive tasks are not purely random systemic noise, but can be partially accounted for by cognitive models of SL as individuals implicitly adapt to stimulus probabilities (Ma & Yu, 2015). An extensive body of research has demonstrated a consistent antagonistic relationship between prefrontal-mediated executive functions and SL. Functional reductions in cognitive control—whether induced by cognitive fatigue (Borragán et al., 2016; Smalle et al., 2022), inhibitory non-invasive brain stimulation targeting the prefrontal cortex (Ambrus et al., 2020; Vékony et al., 2022), or natural transient attentional lapses such as mind wandering (Simor et al., 2025; Vékony, Brezóczki, et al., 2025; Vékony, Farkas, et al., 2025)—actively enhance the implicit extraction of probabilistic sequences. The cognitive states distinguished by RTV variations might therefore be best understood through the lens of the exploration/exploitation framework (Sripada, 2018). Within this paradigm, higher top-down cognitive control drives an "exploitation" state, increasing task focus and decreasing response variability. While this state facilitates better explicit task performance (e.g., faster mean reaction times), it suppresses implicit SL. Conversely, lower cognitive control triggers an "exploration" state characterized by reduced task focus and higher chronometric variability; this degrades immediate task execution but optimally supports the implicit modeling of environmental regularities. Viewed through this lens, our finding that early RTV predicts subsequent learning suggests that elevated chronometric noise acts as a behavioral marker of this relaxed, exploratory cognitive state, unconstrained from prefrontal suppression. This interpretation is corroborated by recent work (Pesthy et al., 2025), which utilized the same experimental paradigm to demonstrate that inhibiting the prefrontal cortex via TMS simultaneously elevates both statistical learning and response variability. Moving forward, further research is necessary to explicitly test the extent to which RTV functions as a reliable marker of neural exploration and, crucially, to clarify the boundary conditions delineating when RTV represents detrimental cognitive decline versus when it acts as a fruitful, adaptive state for learning.

An aim of this study was explicitly methodological, and the comparison across the four RTV indices yields practical recommendations for future work. Earlier studies on variability and learning typically relied on a single summary index, so the question of which aspect of variability actually matters for learning remained open. Our results show that not all RTV measures are equally informative. They differ in what they capture, in how stable they are, and in how well they predict learning. A useful way to organize our findings is to group the four indices by the aspect of the RT distribution they describe. ITRV and Sigma capture ongoing, finer-grained fluctuations in performance, whereas Tau reflects the rightward tail of slow responses usually linked to attentional lapses (Leth-Steensen et al., 2000; Matzke & Wagenmakers, 2009). CV sits somewhere in between, summarizing the overall spread of the distribution relative to the mean. This distinction matters because different parts of the RT distribution reflect distinct cognitive processes (Balota & Yap, 2011; Whelan, 2008), and our data suggest that they also play different roles in predicting learning. ITRV and Sigma were the most robust predictors, whereas Tau was only weakly and inconsistently associated with later learning. The kind of variability that supports SL therefore does not seem to be driven by rare, extreme lapses, but by ongoing, finer-grained fluctuations in performance. Looking at each index in turn helps translate this pattern into concrete recommendations. CV is attractive because it is easy to compute and normalizes the standard deviation by mean RT, which in principle should help separate variability from overall speed. In our data, however, CV showed the lowest test-retest reliability among the four indices and its predictive power replicated only in the larger dataset. We therefore recommend using it with caution, especially in smaller samples. ITRV, which captures the average absolute difference in RT between consecutive trials, emerged as one of the best predictors of later learning, and showed the highest cross-session stability. Because it is simple to compute and does not require distributional assumptions or model fitting, ITRV stands out as a particularly practical choice for studies interested in moment-to-moment fluctuations. The ex-Gaussian parameter Sigma showed a very similar profile. It was the second most stable index across sessions, and it predicted later learning consistently across both datasets. The added value of Sigma is conceptual rather than purely statistical: it isolates the symmetric, Gaussian part of the distribution, and therefore offers a cleaner theoretical interpretation of what is being captured. The downside is that ex-Gaussian fitting is more demanding, both in terms of the amount of data required per participant and in terms of model convergence. In our stability analyses, Sigma and Tau benefited more from additional blocks before their estimates stabilized, whereas ITRV and CV reached a plateau faster. Researchers working with short tasks or few trials per participant should keep this in mind. Tau, finally, behaved differently from the other three indices. Its weak and unstable link with learning is informative rather than disappointing. It suggests that the slow, extreme tail of the RT distribution does not reflect the same kind of variability that supports learning. However, an important caveat is that the metrics demonstrating the highest predictive power were also those that achieved the greatest stability within the initial five blocks. This raises the possibility that Tau and CV are not fundamentally poor predictors of learning; rather, their weak association during early exposure might simply reflect a need for a greater number of trials to generate reliable estimates. Taken together, these comparisons lead to a concrete recommendation. When the goal is to predict learning from early behavior, ITRV and Sigma are the two most useful indices, and we suggest reporting both. ITRV is computationally cheap, robust, and intuitive, and presents the advantage of being temporally resolved. Sigma adds a theoretically grounded decomposition of the RT distribution. CV can be reported as a complementary measure, but it should not be used as the sole index. Tau, while theoretically interesting, appears less suited to predicting this particular form of learning, and its interpretation should be kept separate from that of the other indices.

Despite the conceptual significance of our findings, several limitations outline important avenues for future research. One concerns the size of the effect: early RTV explained roughly 5 to 10 percent of the variance in late stage SL. This figure says as much about the outcome as about the predictfor. Statistical learning estimated within a single ASRT session is a noisy measure, because performance shifts from block to block, and final learning reflects many cognitive, state, and environmental influences that no early marker could capture on its own. We argue that the more telling point is that a short window of early variability predicts later learning at all. The relationship is strong enough to be meaningful, though not yet a tool for predicting how a single individual will learn. A related limitation concerns the specific parameters of the task design. The inter-stimulus interval (ISI), for instance, plays a crucial role in dictating the pacing and cognitive demands of the task, which directly influences the underlying cognitive interpretation of RTV. It is very plausible that altering the temporal constraints of the task could affect the relevance of RTV as a marker of an exploration state. Future studies should systematically manipulate the ISI to evaluate how temporal pressures moderate the predictive relationship between variability and learning. Furthermore, to establish early RTV as a truly domain-general trait marker of learning potential, one must assess its predictive power across a broader battery of paradigms. Testing whether baseline RTV calculated in one context (e.g., a standard sustained attention task) reliably forecasts SL in an entirely different task will be critical for generalizing these findings. Finally, while the present study leveraged traditional summary metrics and ex-Gaussian parameterization, future work should prioritize the integration of generative computational models. Applying frameworks such as drift diffusion models or hidden markov models to trial-by-trial reaction time data would allow researchers to dynamically infer and quantify latent cognitive states, delineating discrete periods of focused task exploitation from exploratory states. Pairing these advanced computational approaches with longitudinal designs—particularly in aging populations where both RTV and learning capacities naturally evolve—would provide a much richer, mechanistic account of exactly when and how behavioral variability facilitates, rather than hinders, the implicit mapping of our environment.

Taken together, these findings carry both a methodological and a theoretical message. On the methodological side, comparing four indices of reaction time variability shows that the choice of measure is not a technical detail: ITRV and Sigma, which track ongoing moment-to-moment fluctuations, reliably predicted later learning and remained stable across sessions, whereas Tau did not. Researchers studying individual differences in learning therefore stand to gain from reporting several variability indices rather than a single summary statistic. On the theoretical side, the results show that variability measured at the very onset of a task, before any structure has been extracted, already anticipates how much implicit statistical learning will follow. Because statistical learning underlies the formation of automatic behaviors, habits, and predictive processing, early variability may offer an early window onto these foundational learning mechanisms. With further work to map its boundary conditions and to extend it across paradigms and populations, this kind of early behavioral signal could eventually inform both educational and clinical settings, wherever the goal is to anticipate, support, or remediate how people learn.

## Data and code availability

The behavioral datasets generated and analyzed during the current study, as well as the custom analysis code, are available in the OSF repository at https://osf.io/87vbs. The experiment was not preregistered.

## Acknowledgement

This work was supported by the French National Research Agency (ANR-24-CE37-5807), the National Brain Research Program (NAP2022-I-2/2022), and the Hungarian Scientific Research Fund (NKFIH ADVANCED153150), all awarded to D.N.; the Spanish Ministry of Science, Innovation and Universities (MICIU), the State Research Agency (AEI), and the European Regional Development Fund (FEDER, UE) through grant PID2024-160183NA-I00 (MICIU/AEI/10.13039/501100011033/FEDER, UE) (T.V.).

We thank Bence Farkas for his valuable advice and comments.

## Declaration on the Use of AI-Assisted Technologies

The authors, who are not native English speakers, used an AI-assisted tool to improve the readability of author-written text. The tool was not used to generate scientific content or ideas. All concepts, analyses, and conclusions are the authors’ own, and the authors take full responsibility for the final manuscript.

## References

Abeles, D., Herszage, J., Shahar, M., & Censor, N. (2023). Initial motor skill performance predicts future performance, but not learning. Scientific Reports, 13(1), 11359. 10.1038/s41598-023-38231-5

Albaugh, M. D., Orr, C., Chaarani, B., Althoff, R. R., Allgaier, N., D’Alberto, N., Hudson, K., Mackey, S., Spechler, P. A., Banaschewski, T., Brühl, R., Bokde, A. L. W., Bromberg, U., Büchel, C., Cattrell, A., Conrod, P. J., Desrivières, S., Flor, H., Frouin, V., … Potter, A. S. (2017). Inattention and reaction time variability are linked to ventromedial prefrontal volume in adolescents. Biological Psychiatry, Attention-Deficit/Hyperactivity Disorder: Predictors of Treatment Response and Comorbidities, 82(9), 660–668. 10.1016/j.biopsych.2017.01.003

Ambrus, G. G., Vékony, T., Janacsek, K., Trimborn, A. B. C., Kovács, G., & Nemeth, D. (2020). When less is more: Enhanced statistical learning of non-adjacent dependencies after disruption of bilateral DLPFC. Journal of Memory and Language, 114, 104144. 10.1016/j.jml.2020.104144

Balota, D. A., & Yap, M. J. (2011). Moving beyond the mean in studies of mental chronometry: The power of response time distributional analyses. Current Directions in Psychological Science, 20(3), 160–166. 10.1177/0963721411408885

Bastian, M., & Sackur, J. (2013). Mind wandering at the fingertips: Automatic parsing of subjective states based on response time variability. Frontiers in Psychology, 4. 10.3389/fpsyg.2013.00573

Booth, T., Dykiert, D., Corley, J., Gow, A. J., Morris, Z., Muñoz Maniega, S., Royle, N. A., del C. Valdés Hernández, M., Starr, J. M., Penke, L., Bastin, M. E., Wardlaw, J. M., & Deary, I. J. (2019). Reaction time variability and brain white matter integrity. Neuropsychology, 33(5), 642–657. 10.1037/neu0000483

Borragán, G., Slama, H., Destrebecqz, A., & Peigneux, P. (2016). Cognitive fatigue facilitates procedural sequence learning. Frontiers in Human Neuroscience, 10. 10.3389/fnhum.2016.00086

Doebler, P., & Scheffler, B. (2016). The relationship of choice reaction time variability and intelligence: A meta-analysis. Learning and Individual Differences, 52, 157–166. 10.1016/j.lindif.2015.02.009

Graveson, J., Bauermeister, S., McKeown, D., & Bunce, D. (2016). Intraindividual reaction time variability, falls, and gait in old age: A systematic review. The Journals of Gerontology Series B: Psychological Sciences and Social Sciences, 71(5), 857–864. 10.1093/geronb/gbv027

He, K., Liang, Y., Abdollahi, F., Fisher Bittmann, M., Kording, K., & Wei, K. (2016). The statistical determinants of the speed of motor learning. PLOS Computational Biology, 12(9), e1005023. 10.1371/journal.pcbi.1005023

Heathcote, A., Popiel, S. J., & Mewhort, D. J. (1991). Analysis of response time distributions: An example using the Stroop task. Psychological Bulletin, 109(2), 340–347. 10.1037/0033-2909.109.2.340

Hervey, A. S., Epstein, J. N., Curry, J. F., Tonev, S., Eugene Arnold, L., Keith Conners, C., Hinshaw, S. P., Swanson, J. M., & Hechtman, L. (2006). Reaction time distribution analysis of neuropsychological performance in an ADHD sample. Child Neuropsychology, 12(2), 125–140. 10.1080/09297040500499081

Howard, J. H., & Howard, D. V. (1997). Age differences in implicit learning of higher order dependencies in serial patterns. Psychology and Aging, 12(4), 634–656. 10.1037/0882-7974.12.4.634

Kaiser, S., Roth, A., Rentrop, M., Friederich, H.-C., Bender, S., & Weisbrod, M. (2008). Intra-individual reaction time variability in schizophrenia, depression and borderline personality disorder. Brain and Cognition, 66(1), 73–82. 10.1016/j.bandc.2007.05.007

Kofler, M. J., Rapport, M. D., Sarver, D. E., Raiker, J. S., Orban, S. A., Friedman, L. M., & Kolomeyer, E. G. (2013). Reaction time variability in ADHD: A meta-analytic review of 319 studies. Clinical Psychology Review, 33(6), 795–811. 10.1016/j.cpr.2013.06.001

Koo, T. K., & Li, M. Y. (2016). A guideline of selecting and reporting intraclass correlation coefficients for reliability research. Journal of Chiropractic Medicine, 15(2), 155–163. 10.1016/j.jcm.2016.02.012

Leth-Steensen, C., King Elbaz, Z., & Douglas, V. I. (2000). Mean response times, variability, and skew in the responding of ADHD children: A response time distributional approach. Acta Psychologica, 104(2), 167–190. 10.1016/S0001-6918(00)00019-6

Leys, C., Ley, C., Klein, O., Bernard, P., & Licata, L. (2013). Detecting outliers: Do not use standard deviation around the mean, use absolute deviation around the median. Journal of Experimental Social Psychology, 49(4), 764–766. 10.1016/j.jesp.2013.03.013

Ma, N., & Yu, A. J. (2015). Statistical learning and adaptive decision-making underlie human response time variability in inhibitory control. Frontiers in Psychology, 6. 10.3389/fpsyg.2015.01046

MacDonald, S. W. S., Nyberg, L., & Bäckman, L. (2006). Intra-individual variability in behavior: Links to brain structure, neurotransmission and neuronal activity. Trends in Neurosciences, 29(8), 474–480. 10.1016/j.tins.2006.06.011

Manly, T., Davison, B., Heutink, J., Galloway, M., & Robertson, I. H. (2000). Not enough time or not enough attention? Speed, error and self-maintained control in the Sustained Attention to Response Test (SART). Clinical Neuropsychological Assessment : An International Journal for Research & Clinical Practice, 3, 167–177.

Matzke, D., & Wagenmakers, E.-J. (2009). Psychological interpretation of the ex-Gaussian and shifted Wald parameters: A diffusion model analysis. Psychonomic Bulletin & Review, 16(5), 798–817. 10.3758/PBR.16.5.798

Oldfield, R. C. (1971). The assessment and analysis of handedness: The Edinburgh inventory. Neuropsychologia, 9(1), 97–113. 10.1016/0028-3932(71)90067-4

Pedraza, F., Vékony, T., Farkas, B. C., Haesebaert, F., Phelipon, R., Mihalecz, I., Janacsek, K., Tillmann, B., Anders, R., Plancher, G., & Nemeth, D. (2025). The interplay between executive functions and updating predictive representations. Scientific Reports, 15(1), 30555. 10.1038/s41598-025-14876-2

Pesthy, O., Pesthy, Z. V., Vékony, T., Janacsek, K., Fabó, D., & Nemeth, D. (2025). Inhibiting the right dorsolateral prefrontal cortex selectively enhances unsupervised statistical learning. bioRxiv. 10.1101/2025.08.08.669288

Rosenberg, M., Noonan, S., DeGutis, J., & Esterman, M. (2013). Sustaining visual attention in the face of distraction: A novel gradual-onset continuous performance task. *Attention, Perception*, & Psychophysics, 75(3), 426–439. 10.3758/s13414-012-0413-x

Simor, P., Vékony, T., Farkas, B. C., Szalárdy, O., Bogdány, T., Brezóczki, B., Csifcsák, G., & Németh, D. (2025). Mind Wandering during Implicit Learning Is Associated with Increased Periodic EEG Activity and Improved Extraction of Hidden Probabilistic Patterns. The Journal of Neuroscience, 45(19), e1421242025. 10.1523/JNEUROSCI.1421-24.2025

Smalle, E. H. M., Daikoku, T., Szmalec, A., Duyck, W., & Möttönen, R. (2022). Unlocking adults’ implicit statistical learning by cognitive depletion. Proceedings of the National Academy of Sciences of the United States of America, 119(2), e2026011119. 10.1073/pnas.2026011119

Song, S., Howard, J. H., & Howard, D. V. (2007). Sleep does not benefit probabilistic motor sequence learning. The Journal of Neuroscience, 27(46), 12475–12483. 10.1523/JNEUROSCI.2062-07.2007

Sripada, C. S. (2018). An Exploration/Exploitation Trade-off Between Mind-Wandering and Goal-Directed Thinking. In K. Christoff & K. C. R. Fox (Eds.), The Oxford Handbook of Spontaneous Thought: Mind-Wandering, Creativity, and Dreaming. Oxford University Press. 10.1093/oxfordhb/9780190464745.013.28

Stark-Inbar, A., Raza, M., Taylor, J. A., & Ivry, R. B. (2017). Individual differences in implicit motor learning: Task specificity in sensorimotor adaptation and sequence learning. Journal of Neurophysiology, 117(1), 412–428. 10.1152/jn.01141.2015

Tamnes, C. K., Fjell, A. M., Westlye, L. T., Østby, Y., & Walhovd, K. B. (2012). Becoming consistent: Developmental reductions in intraindividual variability in reaction time are related to white matter integrity. The Journal of Neuroscience, 32(3), 972–982. 10.1523/JNEUROSCI.4779-11.2012

Vaurio, R. G., Simmonds, D. J., & Mostofsky, S. H. (2009). Increased intra-individual reaction time variability in attention-deficit/hyperactivity disorder across response inhibition tasks with different cognitive demands. Neuropsychologia, 47(12), 2389–2396. 10.1016/j.neuropsychologia.2009.01.022

Vékony, T., Ambrus, G. G., Janacsek, K., & Nemeth, D. (2022). Cautious or causal? Key implicit sequence learning paradigms should not be overlooked when assessing the role of DLPFC (Commentary on Prutean et al.). Cortex, 148, 222–226. 10.1016/j.cortex.2021.10.001

Vékony, T., Brezóczki, B., Csifcsák, G., Németh, D., & Simor, P. (2025). A functional trade-off between executive control and implicit statistical learning is dynamically gated by mind wandering. bioRxiv. 10.1101/2025.08.05.668618

Vékony, T., Farkas, B. C., Brezóczki, B., Mittner, M., Csifcsák, G., Simor, P., & Németh, D. (2025). Mind wandering enhances statistical learning. iScience, 28(2), 111703. 10.1016/j.isci.2024.111703

Verstynen, T., Phillips, J., Braun, E., Workman, B., Schunn, C., & Schneider, W. (2012). Dynamic sensorimotor planning during long-term sequence learning: The role of variability, response chunking and planning errors. PLoS ONE, 7(10), e47336. 10.1371/journal.pone.0047336

Virtanen, P., Gommers, R., Oliphant, T. E., Haberland, M., Reddy, T., Cournapeau, D., Burovski, E., Peterson, P., Weckesser, W., Bright, J., Van Der Walt, S. J., Brett, M., Wilson, J., Millman, K. J., Mayorov, N., Nelson, A. R. J., Jones, E., Kern, R., Larson, E., … Vázquez-Baeza, Y. (2020). SciPy 1.0: Fundamental algorithms for scientific computing in Python. Nature Methods, 17(3), 261–272. 10.1038/s41592-019-0686-2

Whelan, R. (2008). Effective analysis of reaction time data. The Psychological Record, 58(3), 475–482. 10.1007/BF03395630

Wu, H. G., Miyamoto, Y. R., Castro, L. N. G., Ölveczky, B. P., & Smith, M. A. (2014). Temporal structure of motor variability is dynamically regulated and predicts motor learning ability. Nature Neuroscience, 17(2), 312–321. 10.1038/nn.3616

Zanesco, A. P., Denkova, E., & Jha, A. P. (2021). Associations between self-reported spontaneous thought and temporal sequences of EEG microstates. Brain and Cognition, 150, 105696. 10.1016/j.bandc.2021.105696

