## Supplementary Material for "Early reaction time variability predicts implicit statistical learning: a comparison of four variability indices"

### 1. Supplementary methods

### 1.1 Study 1 design

The data for study 1 were drawn from a broader three-day experimental protocol. The primary analyses presented in the current paper utilize data exclusively from Day 1, while data from Day 3 were utilized solely to establish the test-retest stability of the early RTV metrics across days. The study was structured as three consecutive days. On the first day, the experimental task began with three practice blocks of the ASRT task using Sequence 1, where each block contained eighty-five trials. Participants then completed the main learning phase consisting of 25 blocks of the same sequence. In these blocks, the first five trials functioned as a warm-up period. The second day occurred twenty-four hours later. During this session, the sequence differed from the previous day, and participants completed 25 blocks. The third day took place twenty-four hours after the second session and forty-eight hours after the first. This session was designed to test the retrieval and continued learning of both Sequence 1 and Sequence 2. The task was organized into six epochs with three epochs dedicated to each sequence. Each epoch contained five blocks of eighty-five trials, including the five warm-up trials. The order of these sequences was counterbalanced across the participant group.

### 1.2 Quantification of statistical learning as accuracy

Higher values reflect greater implicit statistical learning, indicating that a participant successfully anticipated and responded faster to highly probable sequences. In supplementary analyses, an analogous accuracy-based learning score ($SL_{ACC}$) was computed as the difference in accuracy (mean hit rate) between high- and low-probability triplets:

$$SL_{acc} = Mean\ Accuracy_{High} - \ Mean\ Accuracy_{Low}$$

### 2. Supplementary results

### 2.1 Presence of accuracy-based statistical learning ($SL_{ACC}$)

Before evaluating predictive models, we confirmed the presence of accuracy-based statistical learning during the late blocks (21 to 25). A two-tailed, one-sample *t*-test confirmed that participants responded more accurately to high-probability triplets compared to low-probability triplets in both Dataset 1 ($M$ = 0.013, $SD$ = 0.033, $t(188) = 5.35$, $p < .001$, $d = 0.39$) but not in Dataset 2 ($M$ = 0.010, $SD$ = 0.033, $t(35) = 1.86$, $p = .071$, $d = 0.31$).

### 2.2 Does early RTV predict later $SL_{ACC}$?

To establish whether measures of early RTV predicted SL by the end of the task, a series of simple linear regression analyses was conducted. In Study 1, when SL was indexed by accuracy ($SL_{ACC}$), none of the RTV measures significantly predicted learning outcomes (all $F < 1.12$, all $p > .290$, $R^2 \leq .006$). In Study 2, Sigma emerged as the sole significant positive predictor ($F(1, 34) = 5.26$, $p = .028$, $R^2 = .134$, $b = 0.001$,

$SE = 0.001$, $t = 2.29$). All other early variability measures failed to predict $SL_{ACC}$ (all $F < 1.83$, all $p > .180$, $R^2 \leq .051$). Overall, unlike reaction time-based learning, early RTV is not a reliable predictor of accuracy-based statistical learning.

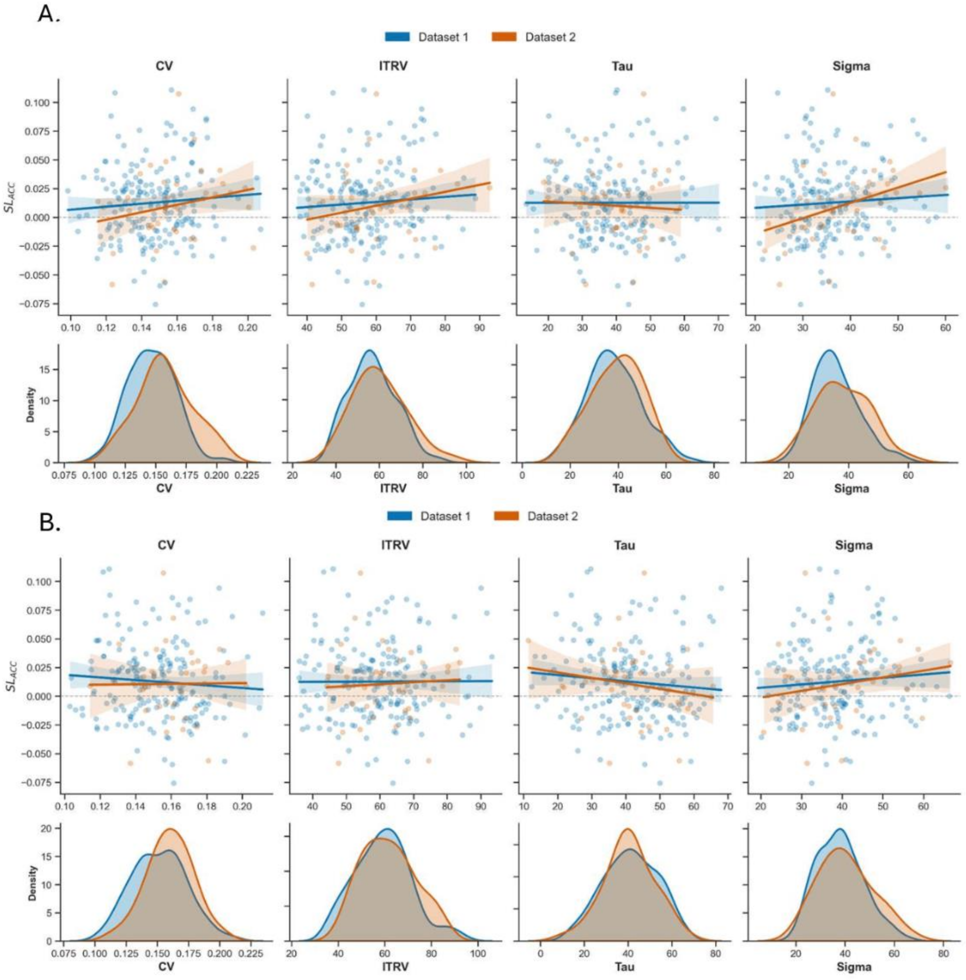


***Supplementary Figure 1.*** *Early RTV does not reliably predict accuracy-based SL ($SL_{ACC}$).* ***(A)*** *RTV calculated during early sequence blocks (Blocks 1–5). Scatterplots and linear regression fits (solid lines with 95% confidence intervals) show the predictive relationship during early sequence exposure. In contrast to the reaction time outcomes in the main text, the regression lines here are predominantly flat. CV, ITRV, and Tau fail to significantly predict accuracy gains in either dataset. The sole exception is Sigma in Dataset 2, which displays a weak but significant positive slope.* ***(B)*** *RTV calculated during sequence-free training blocks. None of the regressions reaches statistical significance, nor is a general stable trend visible across the regressions.*

### 2.3 Cross-Day Prediction of Statistical Learning from RTV

To further investigate the trait-like stability of the relationship between RTV and statistical learning, we conducted supplementary linear regression analyses to test whether early RTV metrics extracted from subsequent experimental sessions (Day 2 and Day 3) could successfully predict initial learning outcomes (Day 1 $SL_{RT}$). Results are summarized in Table 1.

Consistent with the primary analyses, RTV measures from subsequent days remained robust predictors of Day 1 learning. When utilizing Day 2 RTV metrics, CV ($F(1, 182) = 13.95$, $p<.001$, $R^2 = .071$, $b = 86.40$, $SE = 23.14$, $t = 3.73$), ITRV ($F(1, 182) = 11.61$, $p<.001$, $R^2 = .060$, $b = 0.17$, $SE = 0.05$, $t = 3.41$), Sigma ($F(1, 182) = 11.14$, $p = .001$, $R^2 = .058$, $b = 0.24$, $SE = 0.07$, $t = 3.34$), all emerged as significant positive predictors of Day 1 $SL_{RT}$. The parameter Tau showed a marginal, non-significant trend ($F(1, 182) = 3.06$, $p = .082$, $R^2 = .017$, $b = 0.08$, $SE = 0.04$, $t = 1.75$).

A nearly identical pattern was observed when extracting RTV metrics from Day 3. Significant positive associations with Day 1 $SL_{RT}$ were found for CV ($F(1, 179) = 12.65$, $p<.001$, $R^2 = .066$, $b = 78.67$, $SE = 22.12$, $t = 3.56$), ITRV ($F(1, 179) = 11.67$, $p<.001$, $R^2 = .061$, $b = 0.17$, $SE = 0.05$, $t = 3.42$), and Sigma ($F(1, 179) = 11.19$, $p = .001$, $R^2 = .059$, $b = 0.25$, $SE = 0.08$, $t = 3.35$). As in previous models, Tau did not reach statistical significance ($F(1, 179) = 2.71$, $p = .102$, $R^2 = .015$, $b = 0.08$, $SE = 0.05$, $t = 1.65$).

These cross-day predictive models provide strong evidence that early RTV acts as a stable, trait-like cognitive marker that robustly indexes implicit statistical learning capacity, irrespective of the specific day the variability is measured.

**Supplementary Table 1.**

*Linear Regression Results Predicting Day 1 Statistical Learning (*$SL_{RT}$*) from Day 2 and Day 3 Early RTV Metrics*

| **Source Data** | **Predictor** | ***b*** | ***SE*** | **β** | ***t*** | ***p*** | ***F*** | ***R²*** |
|---|---|---|---|---|---|---|---|---|
| *Day 2 RTV* | CV | 86.40 | 23.14 | .27 | 3.73 | < .001 | 13.95*** | .071 |
| *(N = 184)* | ITRV | 0.17 | 0.05 | .24 | 3.41 | < .001 | 11.61*** | .060 |
| | Sigma | 0.24 | 0.07 | .24 | 3.34 | .001 | 11.14** | .058 |
| | Tau | 0.08 | 0.04 | .13 | 1.75 | .082 | 3.06 | .017 |
| *Day 3 RTV* | CV | 78.67 | 22.12 | .26 | 3.56 | < .001 | 12.65*** | .066 |
| *(N = 181)* | ITRV | 0.17 | 0.05 | .25 | 3.42 | < .001 | 11.67*** | .061 |
| | Sigma | 0.25 | 0.08 | .24 | 3.35 | .001 | 11.19** | .059 |
| | Tau | 0.08 | 0.05 | .12 | 1.65 | .102 | 2.71 | .015 |

*Note.* $SL_{RT}$ = Statistical Learning based on Reaction Times; CV = Coefficient of Variation; ITRV = Inter-Trial RTV; β = standardized beta coefficient. Degrees of freedom for Day 2 models are (1, 182) and for Day 3 models are (1, 179). ** $p < .01$; *** $p < .001$.

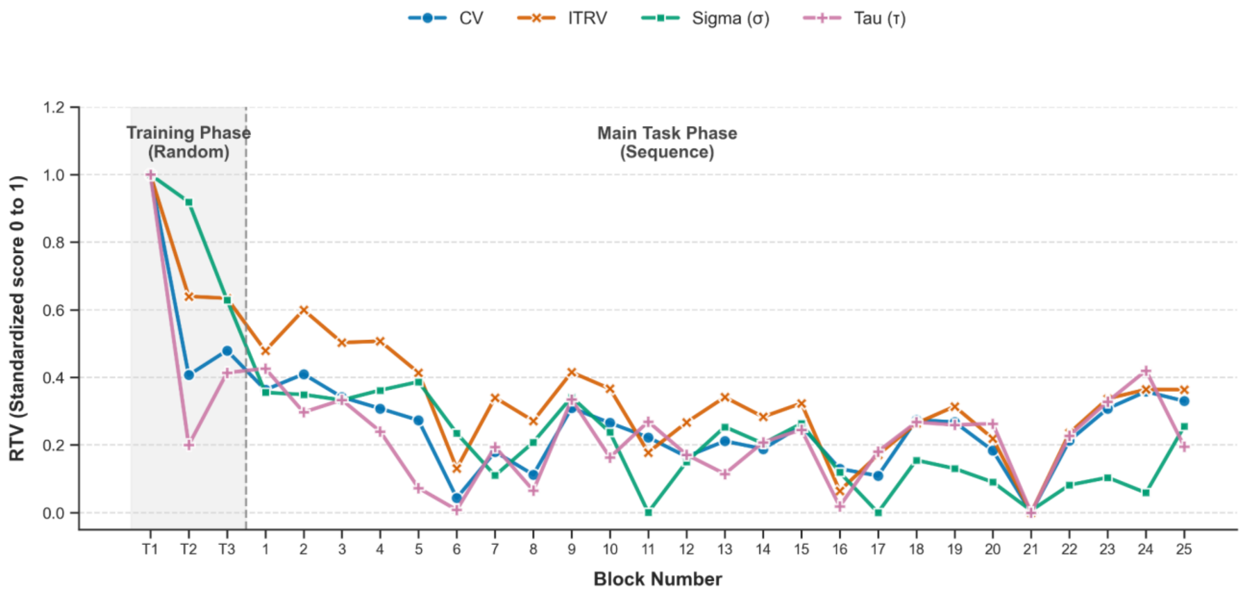


***Supplementary Figure 2***. *Temporal dynamics of RTV throughout the learning paradigm. The line plot illustrates the block-by-block progression of four distinct RTV indices: CV, ITRV, and the ex-Gaussian parameters Sigma and Tau. Data are presented across an initial Training Phase featuring random stimuli (blocks T1–T3, indicated by the shaded background) followed by the Main Task Phase containing the probabilistic sequence (blocks 1–25). To account for differences in raw measurement scales, all metrics were Min-Max standardized, bounding the scores between 0 and 1 to highlight their relative fluctuations over time. Data from Study 1.*

**3. Supplementary Discussion: Divergence between reaction time and accuracy-based learning outcomes**

The analyses revealed a clear dissociation: while early behavioral variability robustly predicted late-stage learning indexed by response speed ($SL_{RT}$), it generally failed to predict learning indexed by accuracy ($SL_{ACC}$). This divergence can be attributed to two primary methodological and cognitive factors.

First, mapping chronometric variability to accuracy outcomes constitutes a cross-domain prediction. RTV metrics are fundamentally speed-based, capturing continuous, moment-to-moment fluctuations in cognitive and motor execution. Consequently, they are conceptually and statistically best aligned with chronometric outcomes. Predicting error rates from speed variability introduces a domain mismatch, as speed and accuracy often reflect distinct strategic components of performance—such as individual differences in speed-accuracy trade-offs—rather than parallel measures of the same underlying cognitive state.

Second, the lack of predictive power for $SL_{ACC}$ is largely driven by the inherent ceiling effects of the ASRT paradigm. In this task, participant accuracy typically starts very high and rapidly reaches a performance plateau (often >90%), with participants making relatively few errors overall. Because of this ceiling effect, the dynamic range of the accuracy metric is severely restricted, stripping away the variance needed to model individual differences. In contrast, reaction times operate on a continuous, wide-ranging scale that remains highly sensitive to implicit probabilistic facilitation long after accuracy has plateaued. Therefore, $SL_{ACC}$ suffers from restricted variance, limiting its utility as a sensitive outcome measure for early chronometric predictors.
